# Heterotrimeric G protein α subunit (GPA1) regulates the response to low-nitrogen stress in *Arabidopsis* by interacting with AtNRT1.4 and AtATG8a

**DOI:** 10.1101/2022.01.27.478073

**Authors:** Mingzhao Luo, Liqin Hu, Weiwei Li, Linhao Ge, Yuhai Qin, Yongbin Zhou, Wensi Tang, Chunxiao Wang, Zhaoshi Xu, Jun Chen, Pierre Delaplace, Youzhi Ma, Ming Chen

**Affiliations:** National Key Facility for Crop Gene Resources and Genetic Improvement, Institute of Crop Sciences, Chinese Academy of Agricultural Sciences, Beijing 100081, China; Beijing Advanced Innovation Center for Food Nutrition and Human Health, Beijing Technology & Business University (BTBU), Beijing 100048, China; University of Liege-GxABT, Agricultural Sciences Department, Plant Sciences and Productions Axis, Plant Biology Laboratory, Passage des Déportés, 2, 5030 Gembloux, Belgium

**Keywords:** *Arabidopsis*, low-nitrogen stress response, heterotrimeric G protein, nitrate transport, autophagy-related protein

## Abstract

Efficient nitrogen absorption and utilization are important factors for higher plants to increase yield and reduce eutrophication (caused by excessive use of nitrogen fertilizers). Heterotrimeric G proteins, including three subunits of α, β, and γ, participate in the pathway regulating nitrogen absorption and utilization in plants. However, the regulatory mechanism remains largely obscured. In this study, our results revealed that the G protein α subunit (AtGPA1) mutant *gpa1-4* was tolerant to low-nitrogen stress in *Arabidopsis*. AtGPA1 was shown to directly interact with a nitrate transporter (AtNRT1.4) and a key autophagy-related protein (AtATG8a) on the plasma membrane using the yeast hybrid system and pull-down assay (*in vitro)* and BiFC assay (*in vivo)*. GUS staining and subcellular localization showed that *AtGPA1* and *AtNRT1.4* were co-expressed in roots and leaf veins and on the plasma membrane. Under low-nitrate conditions, the single mutant *gpa1-4* and *NRT1.4*RNAi plants (*AtNRT1.4*RNA interference plants) and the double mutant *NRT1.4*RNAi/*gpa1-4* plants (*AtNRT1.4*RNA interference plants on a *gpa1-4* background) were healthier than the wild type plants. Moreover, the phenotype of the double mutant *NRT1.4*RNAi/*gpa1-4* plants was closer to that of the *NRT1.4*RNAi plants compared to that of the *gpa1-4* mutants. The results of the nitrate efflux rate assay in roots were consistent with the phenotypic changes under low-nitrogen conditions. These results indicated that AtGPA1 is an upstream factor that regulated the response to low-nitrogen stress through interaction with AtNRT1.4. In addition, we found that transgenic plants overexpressing *AtATG8a* were more tolerant to low-nitrogen stress, and their phenotype was similar to that of *gpa1-4* mutants and double mutant *ATG8aOX/gpa1-4* plants (*AtATG8a* overexpressing plants on a *gpa1-4* mutant background). Further, autophagosome observations were consistent with the phenotypes in mutant plants, indicating that AtGPA1 regulated the response to low-nitrogen stress in *Arabidopsis* plants by affecting the autophagosome assembly. Our findings may provide a new model for improving nitrogen-use efficiency through genetical modification to boost crop yields.

**One sentence summary:** AtGPA1 negative regulates low nitrogen stress response by interaction with a nitrate transporter, AtNRT1.4 and an autophagy-related protein, AtATG8a in *Arabidopsis*.

## INTRODUCTION

In *Arabidopsis*, heterotrimeric G proteins consist of one canonical α subunit (GPA1), one β subunit (AGB1), and at least three γ subunits (AGG), all of which play crucial roles in almost all aspects of growth and development (Pathak et al., 2021). During biotic and abiotic stress, these proteins also control key agronomic traits, including crop yield (Pandey, 2019). Improvement of nitrogen-use efficiency (NUE) is one of the most important goals in agricultural research because it would increase crop yield while reducing eutrophication caused by excessive levels of nitrogen fertilizer in groundwater and rivers (Zhao et al., 2020). Some studies have reported that the intermediaries of the G protein pathway, such as the Gγ subunit encoded by *the dense and erect panicle 1* (*DEP1*) gene that controls panicle branching and erectness, regulate nitrogen absorption and utilization. Further evidence has indicated that *DEP1* can interact with other two G protein subunits, RGA1 (G protein α subunit) and RGB1 (G protein β subunit), to regulate NUE and yield in rice (Huang et al., 2014). Increasing evidence has revealed that RGA1 also regulates the nitrogen-signaling pathway through other genes, such the *bHLH* gene family members, nitrite reductase, *OsCIPK23*, and urea transporters in rice (Pathak et al., 2021). *Setaria* plants overexpressing AGG3 (rice homologs of the *Arabidopsis* group III Gγ subunit) exhibit better growth during early development under conditions of low nitrogen (Kaur et al., 2018). Physiological and molecular evidence in *Arabidopsis* suggests that compared to wild type (WT) plants, the *GPA1* and *GCR1* (*G-protein coupled receptor 1*) double mutant exhibits lower germination and downregulates nitrate transporter-related genes at the transcript level under low-nitrate conditions (Chakraborty et al., 2015; Chakraborty et al., 2019). These studies indicated that the heterotrimeric G protein, especially the Gα subunit, plays an important role in NUE-related regulatory pathways in plants. However, the regulatory mechanism of G proteins in the context of nitrate stress remains largely unknown.

In higher plants, two major gene families involved in nitrate transport—*NRT1* and *NRT2*—and 53 genes have been identified as the *NRT1* family in *Arabidopsis* (Krapp et al., 2014; Wang et al., 2018; Vidal et al., 2020). Among these genes, the most important transporter is *NRT1.1* (formerly *Chl1*), as it not only serves as a dual affinity transporter in response to nitrate but also as a nitrate sensor that regulates nitrate-related pathways in plants (Liu et al., 1999; Liu et al., 2003; Zhang et al., 2020). In response to abscisic acid (ABA) treatment, *NRT1.2* positively regulates seed germination and seedling development by interacting with phospholipase Dα1 (*PLDα1*) (Li et al., 2020). In addition, nitrate is absorbed into the root cells from the external environment, and it can be transported to various types of cells and different part of tissues in whole plants. Both *NRT1.5* and *NRT1.8* are important for this nitrate transport process. For example, *NRT1.5* was the first transporter that regulated the long-distance translocation of nitrate from root to shoot; further, in-situ hybridization showed the expression of *NRT1.5* in the pericycle cells of the central vascular system. This suggested that *NRT1.5* is also responsible for nitrate loading in xylem vessels. In contrast, the function of *NRT1.8* may differ from that of *NRT1.5* with respect to nitrate distribution, wherein it can serve as a nitrate-unloading transporter in xylem vessels (Lin et al, 2008; Li et al., 2010; Meng et al., 2016; Watanabe et al., 2020). The *NRT1.6* gene is only expressed in reproductive tissue, and its expression increases after pollination; additionally, it can promote seed development by increasing nitrate accumulation in the developing embryo (Almagro et al., 2008; O’Brien et al., 2016). Although *NRT1.7* is only expressed in the phloem tissue of the minor veins in the distal leaf part, it shows higher expression in old leaves than in young leaves. Moreover, *NRT1.7* plays a key role in nitrate remobilization from source leaves (older leaves) to nitrate-demanding tissues (younger leaves) to adapt to low nitrogen levels (Fan et al., 2009; Chen et al., 2020). *NRT1.9* is mainly expressed in the companion cells of root vascular tissues. Functional analysis showed that *NRT1.9* can load nitrate into the phloem of root and mediate nitrate distribution from root to shoot tissues (Wang et al., 2011). *NRT1.11* and *NRT1.12* are functionally redundant, both of which are expressed in the petiole and major vein of leaves. Additionally, they are responsible for loading nitrate into the phloem from the xylem and controlling nitrate transfer into younger leaves (Hsu et al., 2013). Recently, a new low-affinity nitrate transporter, *NRT1.13*, was found to modulate the flowering time, and it can influence the node number and basal branch under low-nitrate conditions (Chen et al., 2021). *NRT1.4* was first revealed to store nitrate in the leaf petiole; further, the nitrate content in the petioles of mutant *nrt1.4* plants is only half of that in the petioles of WT plants (Chiu et al., 2004; Morales et al., 2021).

Autophagy, a highly conserved mechanism that degrades and recycles intracellular materials to promote survival under different external environment stresses, exists extensively in animals and plants (Fan et al., 2020; Melino et al., 2021). In plants, autophagy plays a crucial role in nutrient recycling under conditions of nitrogen or carbon starvation (Chen et al., 2019; Zhen et al., 2019a; Zhen et al., 2019b; Zhen et al., 2021). Most autophagy-related genes (ATGs) are upregulated in response to nitrogen starvation, and in *Arabidopsis*, mutations in key autophagy-related genes, such as *atg5, atg7, atg10*, and *atg13a*/*atg13b*, increase the sensitivity to low-nitrate stress, accelerate senescence, and enable the supply of sufficient nitrate (Zhao et al., 2019; Havé et al., 2019). Some other *ATG* mutants in *Arabidopsis* and maize significantly influence the seed traits by regulating nitrate remobilization; moreover, 50% of the remobilized nitrogen is derived from autophagy in *Arabidopsis* (Zhou et al., 2013; Pottier et al., 2014). To date, over 40 ATGs have been demonstrated to have conserved functions from prokaryotes to eukaryotes (Marshall et al., 2018). Unlike the ubiquitin-proteasome system, autophagy can breakdown various macromolecules to digest organelles through autophagosomes. Multiple highly conserved ATGs are involved in autophagosome formation. ATG8 and ATG12 belong to the ubiquitin-like conjugation system, of which ATG8 is crucial for membrane scaffold formation, elongation, and expansion during autophagosome generation (Marshall et al., 2019; Wesch et al., 2020; Johansen et al., 2020). Overexpression of *AtATG8f* and *GmATG8c* enhances tolerance to both nitrogen and carbon starvation in *Arabidopsis* (Jacomin et al., 2020). Moreover, interactions with *ATG8* are important for regulating autophagy. For instance, constitutively stressed 1 (*AtCOST1*) restricts autophagosome formation to promote plant growth by interacting with *AtATG8e* directly under normal conditions; however, drought stress induces the degradation of *COST1* to release *ATG8* and promotes autophagosome formation to improve drought resistance (Bao et al., 2020). In addition, a ubiquitous protein in the plant kingdom, AtNBR1, directly binds to AtATG8 to regulate the accumulation of insoluble and detergent-resistant proteins (Zhou et al., 2013). In this study, we found that the *gpa1-4* mutant had a high tolerance to low-nitrogen stress in *Arabidopsis*. We used AtGPA1 as bait to screen interaction proteins, and found that AtGPA1 could interact with AtNRT1.4 (a low-affinity nitrate transporter) and AtATG8a (a key autophagosome-related protein) in *Arabidopsis*. Biochemical and genetic analyses showed that AtGPA1 interacted with AtNRT1.4 and AtATG8a to regulate nitrogen absorption, nitrogen transport, and autophagy assembly. These processes restricted the flexibility to low-nitrogen stress in *Arabidopsis*.

## RESULTS

### *AtGPA1* Mutant *gpa1-4* had Higher Tolerance to Low-Nitrogen Stress than the WT

To study the function of *AtGPA1* under low-nitrogen stress, we first identified the homozygote mutant, *gpa1-4* (Supplemental Fig. S1C). To examine the phenotype of *gpa1-4*, we planted WT and *gpa1-4* under normal and low-nitrogen conditions through a hydroponic experiment (Supplemental Fig. S1A, B). The results showed that the *gpa1-4* mutant had significantly higher leaf width than WT under normal and low-nitrogen conditions, although leaf length was not significantly different between the groups (Supplemental Fig. S1A, B, D, E). In addition, we planted mutant *gpa1-4* on a solid medium with low nitrate content (see Supplemental Table S1 for medium composition). We found that the fresh weight, chlorophyll content, total root surface area, and total root length were higher in *gpa1-4* mutants than in WT under low-nitrogen conditions, although no differences were observed under normal conditions (Fig. 1A–F). We also examined the subcellular structure of *gpa1-4* and WT under normal and low-nitrate conditions. This examination revealed that *gpa1-4* mutants had healthier chloroplasts than the WT, and the difference was more significant under low-nitrate conditions than under normal conditions (Fig. 1G–J). These phenotypic results suggested that AtGPA1 negatively regulates tolerance to low-nitrogen stress in *Arabidopsis*.

**Figure 1.**
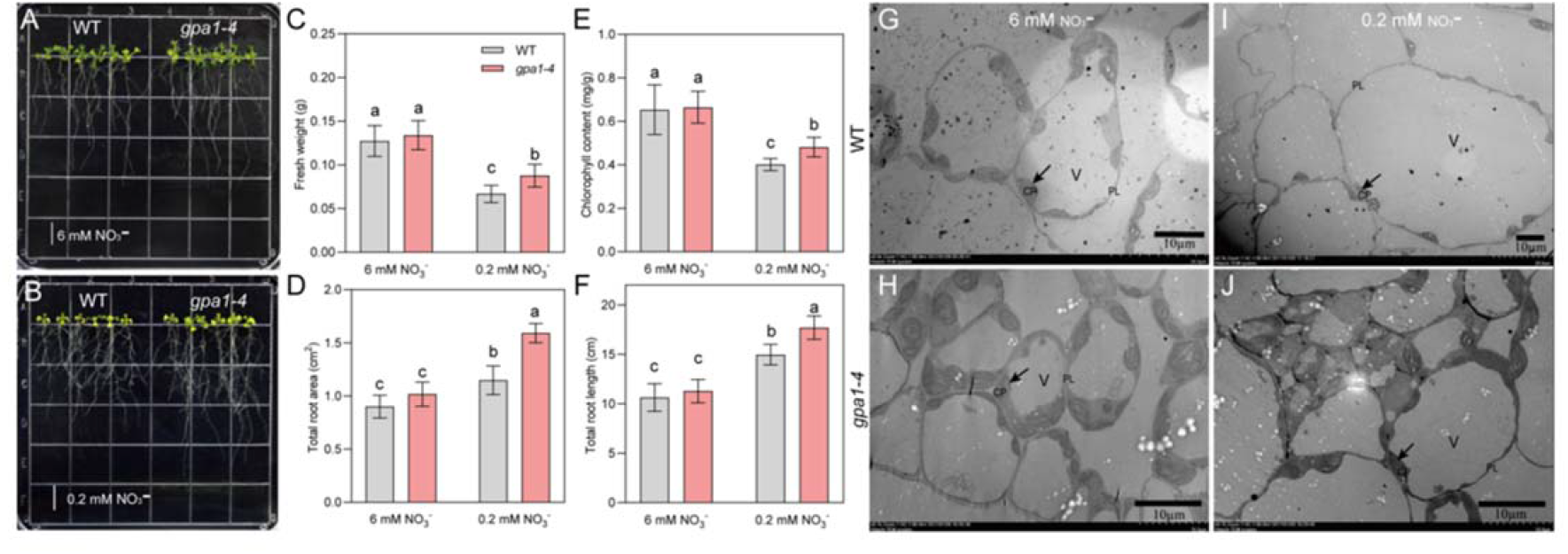
Phenotype of *gpa1-4* mutants and the wild type (WT) under low-nitrate conditions. A, Phenotype of *gpa1-4* and WT grown on normal-nitrogen medium (6 mM NO_3_^−^). B, Phenotype of *gpa1-4* and WT grown on low-nitrogen medium (0.2 mM NO_3_ ^−^). Scale bars = 1 cm. C, D, E and F, The *gpa1-4* mutants have higher fresh weight (C), total root surface area (D), chlorophyll content (E), and total root length (F) under conditions of nitrate starvation (0.2 mM NO_3_^−^). The *gpa1-4* and WT plants had no significant growth difference in the normal-nitrogen medium. G and H, Subcellular structure of mesophyll cells in *gpa1-4* and WT under normal conditions. I and J, Subcellular structure of mesophyll cells in *gpa1-4* and WT under low-nitrate conditions. Black arrows indicate chloroplasts in mesophyll cells. V: vacuole. CP: chloroplast. PL: plasmalemma. Scale bars = 10 μm. All data represent means ± SD (standard deviation) (n = 6). Different letters above the bars represent significant differences between mutants and WT (*P* < 0.05 according to one-way analysis of variance (ANOVA) followed by Duncan’s multiple range test).

### AtGPA1 Interacted with AtNRT1.4 and AtATG8a on the Plasma Membrane

To identify AtGPA1 interaction proteins in *Arabidopsis*, we screened yeast cDNA libraries constructed with total RNA from plants treated with low-nitrogen conditions. We observed that AtGPA1 interacted with AtNRT1.4 (a nitrate transport protein) and AtATG8a (a key autophagosome-related protein). Because AtGPA1 is located on the plasma membrane, we used the DUAL membrane interaction protein screening system to identify interactions in yeast cells. The positive controls were the membrane interacting proteins pNubG-Fe65 and pTSU2-APPs (Fig. 2A). The screening results showed that the yeast cells with two pairs of interacting proteins, AtGPA1 and AtNRT1.4, and AtGPA1 and AtATG8a, could grow on the medium lacking four nutrients, whereas the negative control could not grow on this medium. This suggests that AtGPA1 can interact with both AtNRT1.4 and AtATG8a in yeast cells (Fig. 2A). We confirmed the direct interaction between AtGPA1 and AtNRT1.4 using glutathione S-transferase (GST) pull-down assays. The results showed that the fusion protein, GST-AtGPA1, could pull down the fusion protein, His-AtNRT1.4, from the solution. Thus, His-AtNRT1.4 could be detected with anti-His antibody upon elution from an affinity chromatography column (Fig. 2B). Furthermore, we identified these interactions in plant cells using BiFC assay. Vectors containing the interacting proteins (AtGPA1 and AtNRT1.4 or AtGPA1 and AtATG8a) were transformed into the mesophyll cells protoplast of *Arabidopsis* through *Agrobacterium tumefaciens*-mediated transformation. The results showed that yellow fluorescence signals were only detected on the plasma membrane when both nYFP-AtGPA1 and cYFP-AtNRT1.4, or both nYFP-AtGPA1 and cYFP-AtATG8a vectors were co-expressed in *Arabidopsis* cells. In contrast, no YFP signals were observed in the negative control (Fig. 2C). These results showed that AtGPA1 directly interacted with AtNRT1.4 and AtATG8a on the plasma membrane.

**Figure 2.**
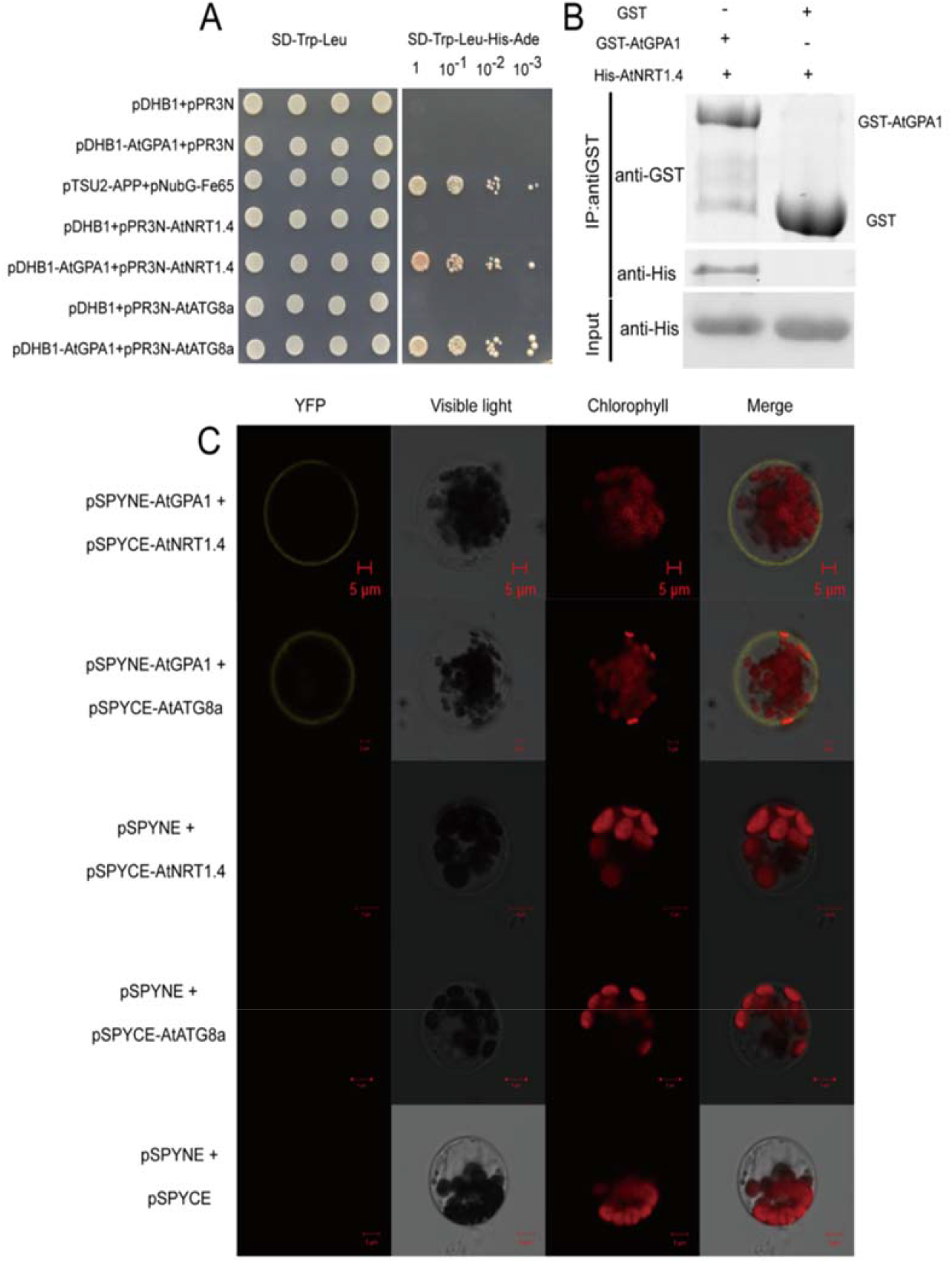
AtGPA1 interacts with AtNRT1.4 and AtATG8a. A, AtGPA1 can interact with AtNRT1.4 in yeast cells. The interaction was assessed according to the growth status of yeast cells. The positive controls were membrane interacting proteins, prey pNubG-Fe65 (expressing the cytosolic protein, Fe65) and the bait pTSU2-APP (expressing the type I integral membrane protein, APP) in the DUAL membrane functional assay. The empty vectors, pDHB1 and pPR3N, were used as negative controls. B, AtGPA1 interacts with AtNRT1.4, as identified by the GST pull-down assay. The “–” or “+” symbols represent the absence or presence of corresponding proteins, respectively. C, AtGPA1 interacts with AtNRT1.4 and AtATG8a on plasma membrane, as identified by the BiFC assay. Left to right: YFP (yellow) fluorescence, a bright-field image, chlorophyll (red) fluorescence, and overlap of the YFP (yellow) and chlorophyll (red) fluorescence. The empty vectors, pSPYNE and pSPYCE, were used as negative controls. Scale bars = 5 μm.

### *AtGPA1* and *AtNRT1.4* showed Similar Tissue-specific Expression Patterns and Subcellular Localization

To analyze the tissue expression patterns of *AtGPA1* and *AtNRT1.4*, we used *GUS* as a reporter gene. The promoters of *AtGPA1* and *AtNRT1.4* were fused with the GUS gene to generate the respective vectors, *proAtGPA1:GUS* and *proAtNRT1.4:GUS*, which were then transformed into *Arabidopsis* via *Agrobacterium*-mediated transformation. The results suggested that both *AtGPA1* and *AtNRT1.4* had high expression levels in the roots and leaf veins, whereas in stems, *AtGPA1* was highly expressed and *AtNRT1.4* was not expressed (Fig. 3A–F). These expression patterns suggested that *AtGPA1* functions in roots, leaf veins, and stems, whereas *AtNRT1.4* mainly functions in roots and leaf veins. To analyze the expression profile of *AtGPA1* and *AtNRT1.4* at the subcellular level, fusion vectors (*AtGPA1-GFP* or *AtNRT1.4-GFP*) were transiently transformed into *Arabidopsis* mesophyll protoplasts. The merged field image obtained using confocal microscopy revealed that both proteins were expressed in the plasma membrane, whereas the positive control was detected in almost all subcellular structures (Fig. 3G–I). These results indicated that *AtGPA1* and *AtNRT1.4* were co-localized in the plasma membrane. These co-localization results of *AtGPA1* and *AtNRT1.4* were consistent with their BiFC results.

**Figure 3.**
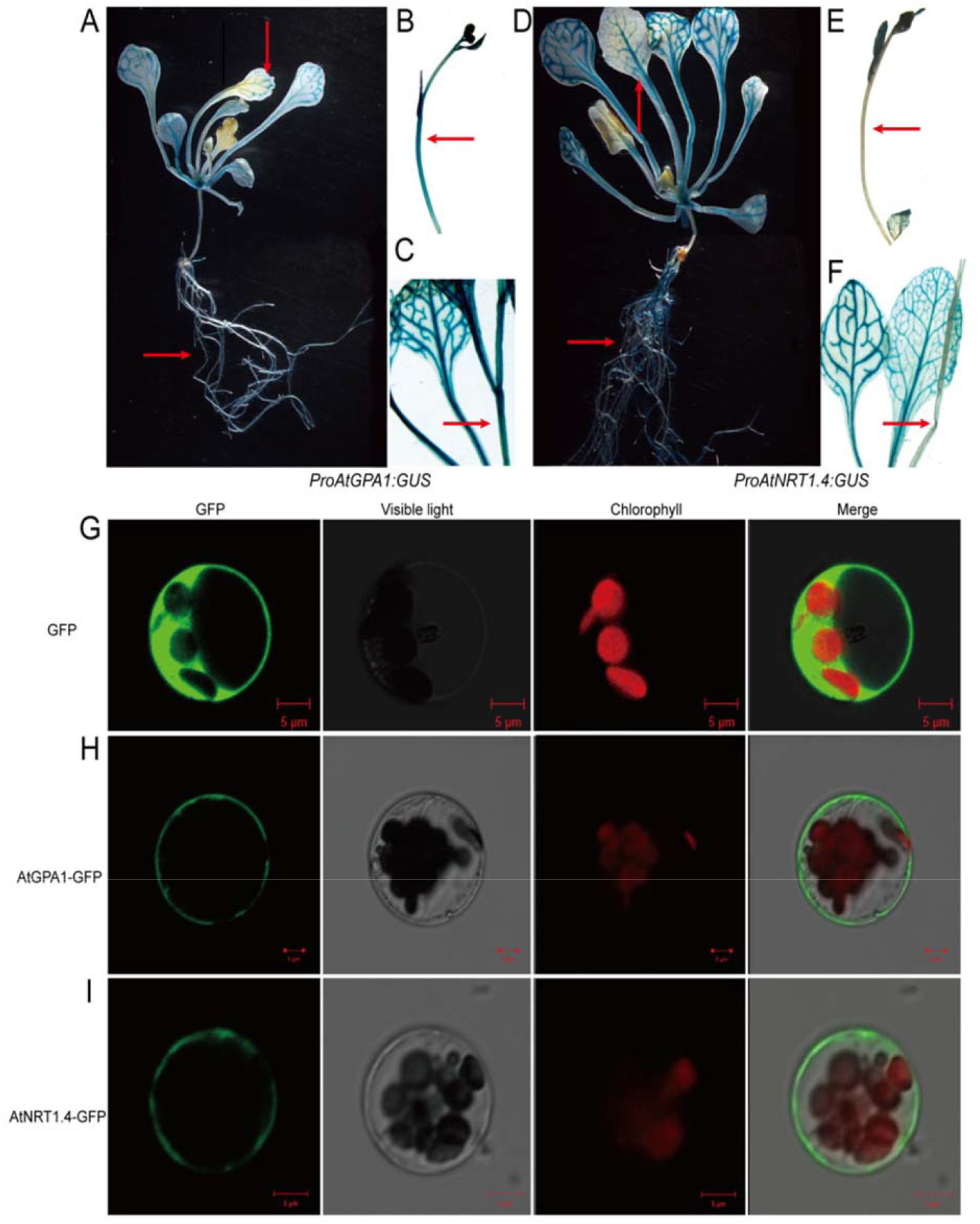
Tissue-specific expression and subcellular localization of *AtGPA1* and *AtNRT1.4*. A, and D, Tissue-specific expression of *AtGPA1* and *AtNRT1.4* in *Arabidopsis* plants. *AtGPA1* and *AtNRT1.4* are co-expressed mainly in the root and leaf veins. B, C, E, and F, *AtGPA1* is highly expressed in the stems, whereas *AtNRT1.4* is not expressed in the stems. Red arrows indicate differential expression between *AtGPA1* and *AtNRT1.4* in the stems. G, Results of a subcellular localization analysis using a GFP control. H, and I, Microscopy images of *Arabidopsis* protoplast expressing the AtGPA1-GFP and AtNRT1.4-GFP fusion genes. GFP fluorescence is detected in the plasma membrane. Left to right: GFP (green) fluorescence, a bright-field image, chlorophyll (red) fluorescence, and overlap of the GFP (green) and chlorophyll (red) fluorescence. Scale bars = 5 μm.

### *AtGPA1* Regulated Tolerance to Low-Nitrogen Stress through *AtNRT1.4*

To confirm that *AtNRT1.4* is involved in tolerance to low-nitrogen stress, we generated *NRT1.4*RNAi mutant lines using the vacuum infiltration method in *Arabidopsis*. The expression of *AtNRT1.4* was inhibited in *NRT1.4*RNAi lines #12, #14, and #38 (Supplemental Fig. S2B). We cultured these lines and WT under normal and low-nitrogen conditions for 7 d. The results showed that in the low-nitrogen medium, *NRT1.4*RNAi lines *#*12, #14, and #38 had significantly higher total root length, total root surface, fresh weight, and chlorophyll content than the WT. However, in the normal-nitrogen medium, there was no significant difference between *NRT1.4*RNAi lines and WT (Fig. 4, Supplemental Fig. S3A–F). These results indicated that *AtNRT1.4* negatively regulated the response to nitrate starvation in *Arabidopsis*, which is similar to the response observed in the *AtGPA1* mutant *gpa1-4* (Fig. 4).

**Figure 4.**
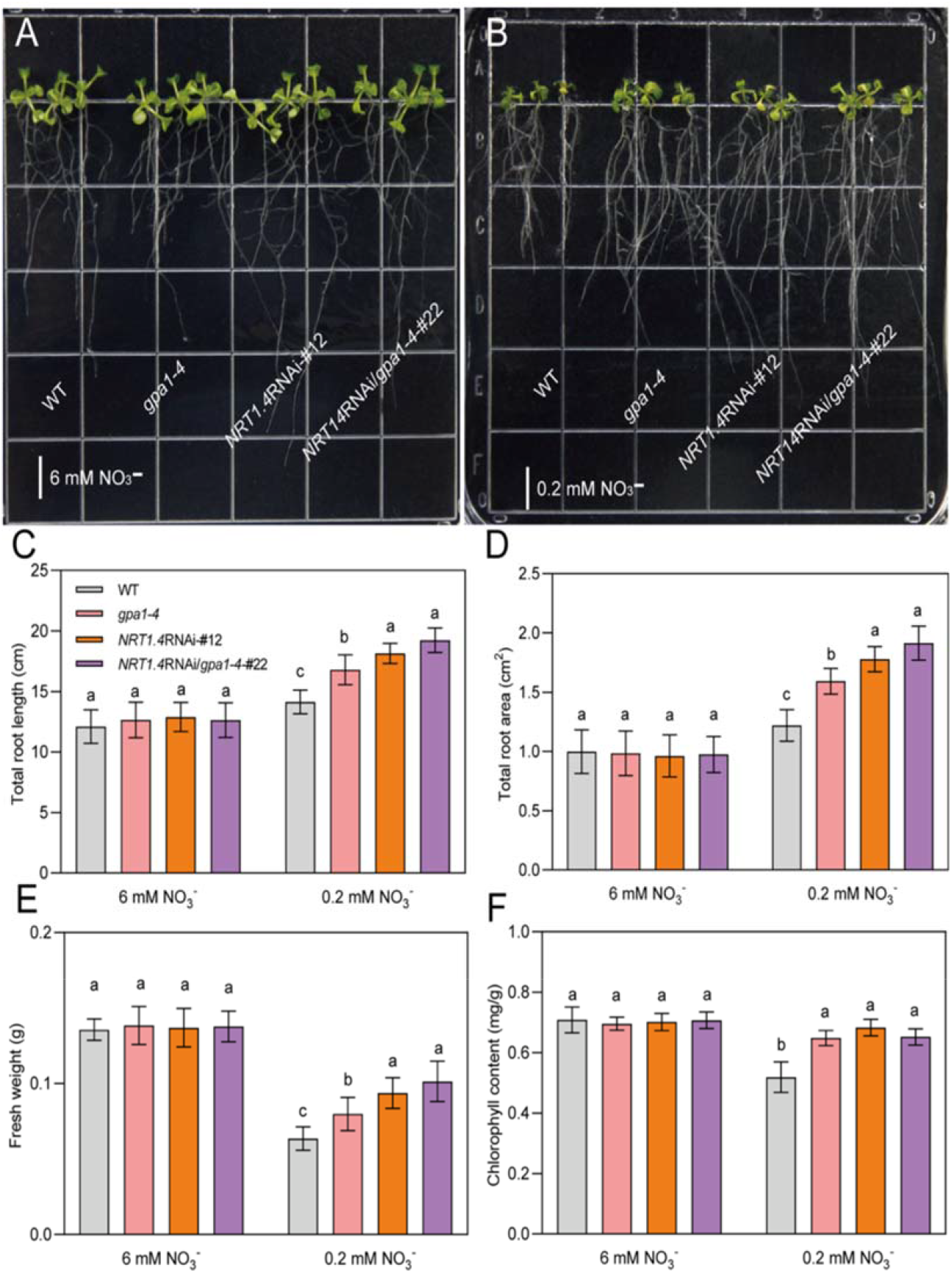
Phenotype of the single mutants, *gpa1-4* and *NRT1.4*RNAi, and the double mutant, *NRT1.4*RNAi/*gpa1-4*, under low-nitrogen conditions. A and B, Phenotype of WT, *gpa1-4, NRT1.4*RNAi #38, and *NRT1.4*RNAi*/gpa1-4* #22 grown on normal-nitrogen medium (6 mM NO_3_^−^) and low-nitrogen medium (0.2 mM NO_3_^−^). C, D, E, and F, Double mutant *NRT1.4*RNAi*/gpa1-4* lines have significantly higher total root length, total root surface, fresh weight, and chlorophyll content than WT plants grown under low-nitrogen conditions; the phenotype of the double mutant is more similar to that of the *NRT1.4*RNAi mutant than to the phenotype of the *gpa1-4* mutant. All data are represented as mean ± standard deviation (SD) (n = 6). Different letters above bars represent significant differences by difference analysis between mutants and WT (*P* < 0.05 according to one-way ANOVA followed by Ducan’s multiple range test). Scale bars = 1 cm.

To verify the epistatic relationship between *AtGPA1* and *AtNRT1.4*, we hybridized mutant *NRT1.4*RNAi plants (#12 and #38) and the *gpa1-4* mutant to generate the *NRT1.4*RNAi*/gpa1-4* double mutant line. *NRT1.4*RNAi*/gpa1-4* double mutant lines #22 and #25 were the hybrid offspring of *NRT1.4*RNAi #12 and *gpa1-4*, whereas *NRT1.4*RNAi*/gpa1-4* double mutant line #32 was the hybrid offspring of *NRT1.4*RNAi #38 and *gpa1-4*. We detected *AtNRT1.4* expression in all double mutant plants, and found that the expression of *AtNRT1.4* was inhibited in three double mutant lines: #22, #25, and #32 (Supplemental Fig. S2B). Phenotypic analysis showed that in the low-nitrogen medium, the double mutant *NRT1.4*RNAi*/gpa1-4* lines had significantly higher total root length, total root surface, fresh weight, and chlorophyll content than the WT (Supplemental Fig. S4A–F). The single mutants, *gpa1-4* and *NRT1.4*RNAi mutant line #12, and double mutant *NRT1.4*RNAi*/gpa1-4* #22 were also grown on low-nitrogen medium. Phenotype analysis of these lines showed that double mutant #22 had higher total root length, total root surface, fresh weight, and chlorophyll content compared to the WT. Under low-nitrogen conditions, the total root length, total root surface, and fresh weight of *NRT1.4*RNAi*/gpa1-4* #22 mutants were more similar to those of *NRT1.4*RNAi single mutants than to those of *gpa1-4* mutants (Fig. 4A–F). This suggested that *AtGPA1* and *AtNRT1.4* were involved in the same pathway to regulate the response to nitrogen. Moreover, *AtGPA1* was upstream of *AtNRT1.4* in the pathway.

### *AtGPA1* Negatively Regulated Nitrate Uptake in Root Hair and Root Tip Zones through *AtNRT1.4*

We used the scanning ion-selective electrode technique (SIET) to analyze the nitrate flow state at the root tip and root hair zones under normal and low-nitrate conditions. The results indicated that the nitrate efflux rate of the single mutant (*gpa1-4* and *NRT1.4*RNAi) and double mutant (*NRT1.4*RNAi*/gpa1-4*) was significantly lower than that of the WT in both the root tip zone and the root hair zone under normal or low-nitrogen conditions (Fig. 5A–D). Under low-nitrate conditions, the nitrate efflux rate of the WT in the root tip zone was 63.45 pmol·cm^−2^·s^−1^, and the average NO_3_^−^ efflux rates of the *gpa1-4, NRT1.4*RNAi, and *NRT1.4*RNAi*/gpa1-4* mutants were 44.97, 25.36, and 13.82 pmol·cm^−2^·s^−1^, respectively (Fig. 5E). This indicated that the nitrate efflux rate of all mutants was lower than that of the WT under low-nitrate conditions in the root tip zone. In the root hair zone, the average NO_3_^−^ efflux rates of the WT and the *gpa1-4, NRT1.4*RNAi, and *NRT1.4*RNAi*/gpa1-4* mutants were 81.95, 54.73, 20.24, and 7.3 pmol·cm^−2^·s^−1^, respectively, under low-nitrogen conditions (Fig. 5F). These results were similar to those observed in the root tip zone. In addition, we found that the nitrate efflux rates of the double mutant (*NRT1.4*RNAi/*gpa1-4*) in the root tip or root hair zones were closer to those of the *NRT1.4*RNAi mutant under low-nitrogen conditions than to those of the *gpa1-4* mutant. These results indicated that both the single mutants (*gpa1-4* and *NRT1.4*RNAi) and the double mutant (*NRT1.4*RNAi*/gpa1-4*) accumulated more nitrate in their roots than the WT under low-nitrogen conditions. Moreover, nitrate accumulation in the double mutant was closer to that of *NRT1.4*RNAi than to that of *gpa1-4*. The nitrate flux rates of all mutants were consistent with their phenotype observations under low-nitrogen conditions (Fig. 4).

**Figure 5.**
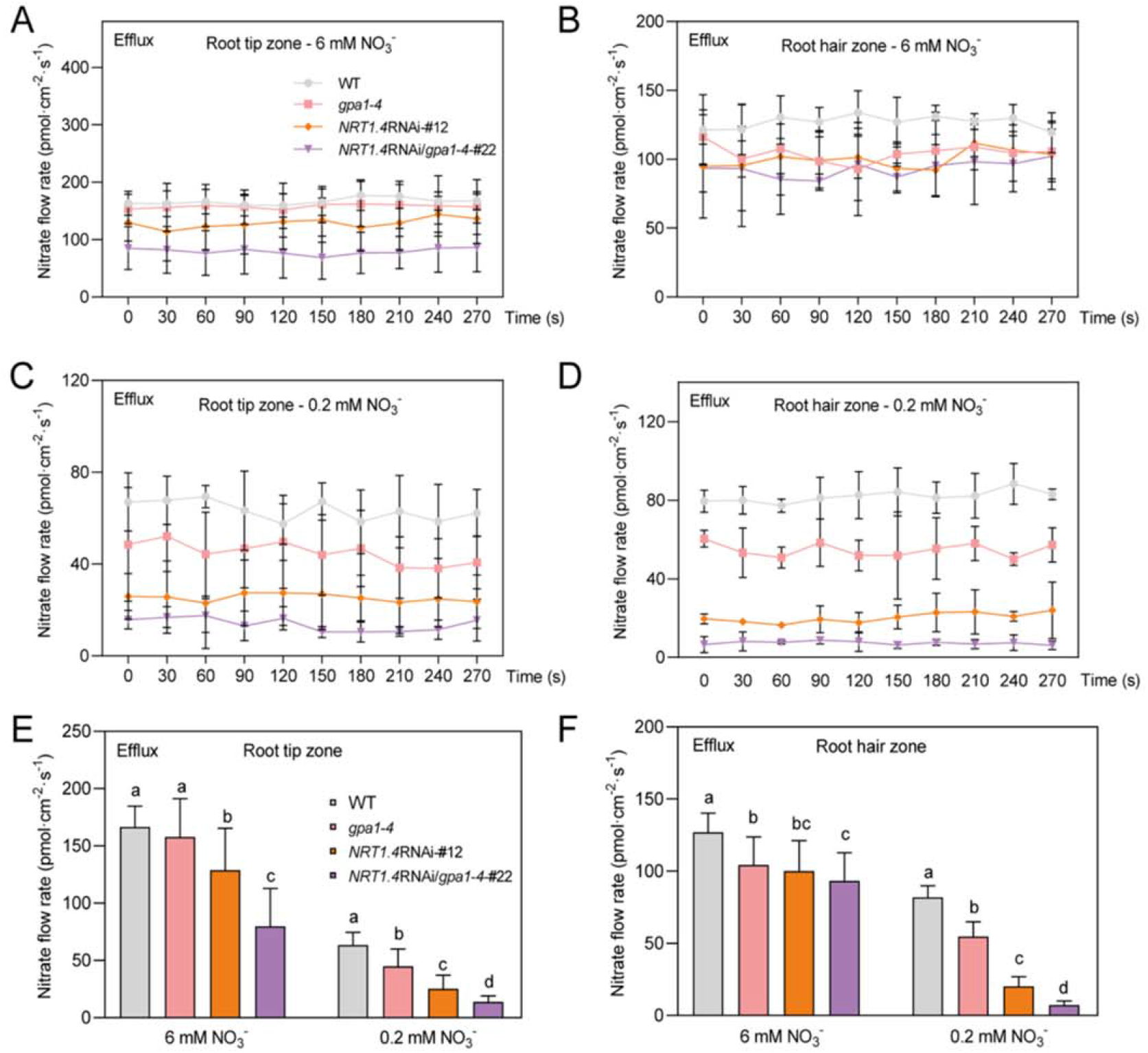
Nitrate efflux rate of WT and different mutants in the root tip and root hair zones under normal or low-nitrate conditions. A and C, Nitrate efflux flow rates in the root tip zone of plants grown in liquid medium containing 6 mM or 0.2 mM NO_3_^−^. B and D, Nitrate efflux flow rates in the root hair zone of plants grown in liquid medium containing 6 mM or 0.2 mM NO_3_ ^−^. E, Mean values of all nitrate efflux flow rates in the root tip zone from (A) and (C). F, Mean values of all nitrate efflux flow rates in the root hair zone from (B) and (D). All data are represented as mean ± SD (n= 6). Different letters above the bars represent significant differences between the mutants and WT (*P* < 0.05 according to one-way ANOVA followed by Ducan’s multiple range test).

### *AtGPA1* Regulated the Response to Low-Nitrogen Stress through *AtATG8a*

Because the functions of different members of ATG8-like proteins are redundant (Bu et al., 2020; Jacomin et al., 2020), we overexpressed *AtATG8a* in the WT background to generate transgenic lines, such as *ATG8aOX* #3, #5, and #7, to identify the functions of *AtATG8a* under low-nitrogen conditions (Supplemental Fig. S2C).

Under low-nitrogen conditions, transgenic lines with overexpressed *AtATG8a* had significantly higher total root length, total root surface, fresh weight, and chlorophyll content than the WT. In contrast, there were no significant differences between the groups under normal-nitrogen conditions (Supplemental Fig. S5A–F). *AtATG8a* was overexpressed in the *gpa1-4* mutant background to generate the double mutant *ATG8aOX/gpa1-4* by crossing the *AtATG8a* overexpression transgenic lines with *gpa1-4. ATG8aOX/gpa1-4* double mutant lines #3 and #6 were the hybrid offspring of *ATG8aOX* #3 and *gpa1-4*, whereas *ATG8aOX/gpa1-4* double mutant line #17 was the hybrid offspring of *ATG8aOX* #7 and *gpa1-4* (Supplemental Fig. S2C). We found that all double mutant plants had higher total root length, total root surface, fresh weight, and chlorophyll content than the WT under low-nitrogen conditions (Supplemental Fig. S6A–F). The single mutants (*gpa1-4* and *ATG8aOX* transgenic lines) and the double mutant (*ATG8aOX/gpa1-4*) also had higher total root length, total root surface, fresh weight, and chlorophyll content compared to the WT (Fig. 6). In addition, we observed autophagosome accumulation in different genotypes under low-nitrogen conditions. The *AtATG8a* gene was fused with the *GFP* gene in the WT or *gpa1-4* mutant background. In the WT, low-nitrogen starvation induced more autophagosome accumulation than normal conditions. Under low-nitrogen conditions, autophagosome accumulation increased to a greater degree in the *AtATG8a*-overexpressing transgenic plants on the *gpa1-4* mutant background than in *AtATG8a*-overexpressing transgenic plants on the WT background (Fig. 7A–C). In *ATG8aOX* transgenic plants, the phenotype of autophagosome accumulation under low-nitrogen conditions was consistent with the phenotypic changes in the roots and shoots, indicating that *AtGPA1* and *AtATG8a* are involved in the same pathway that regulates the response to low-nitrogen stress in *Arabidopsis*.

**Figure 6.**
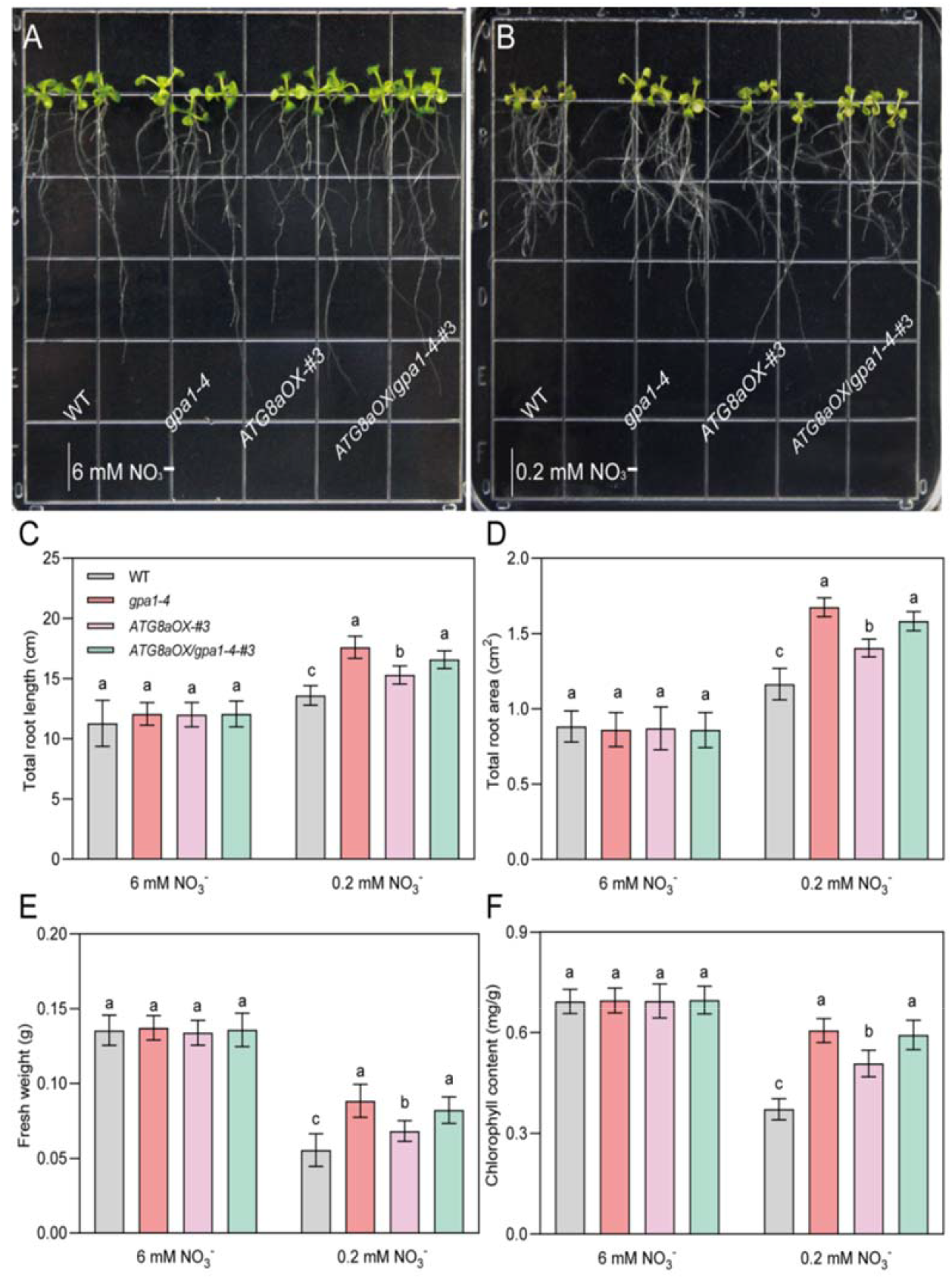
*AtATG8a* overexpression in WT and *gpa1-4* backgrounds increases tolerance to low-nitrate stress. A and B, Phenotypes of WT, *gpa1-4, ATG8aOX* overexpression line #3, and double mutant *ATG8aOX/gpa1-4* line #3 grown on normal-nitrogen medium (6 mM NO_3_^−^) and low-nitrogen medium (0.2 mM NO_3_^−^). C, D, E, and F, *ATG8aOX* overexpression line *#3* has significantly higher total root length, total root surface, fresh weight, and chlorophyll content than WT plants grown on low-nitrogen medium. The *gpa1-4* mutants and *ATG8aOX/gpa1-4* #3 double mutants have better root development and higher fresh weight than WT plants grown under low-nitrate conditions. All data are represented as mean ± SD (n = 6). Different letters above the bars represent significant differences between the mutants and WT (*P* < 0.05 according to one-way ANOVA followed by Ducan’s multiple range test). Scale bars = 1 cm.

**Figure 7.**
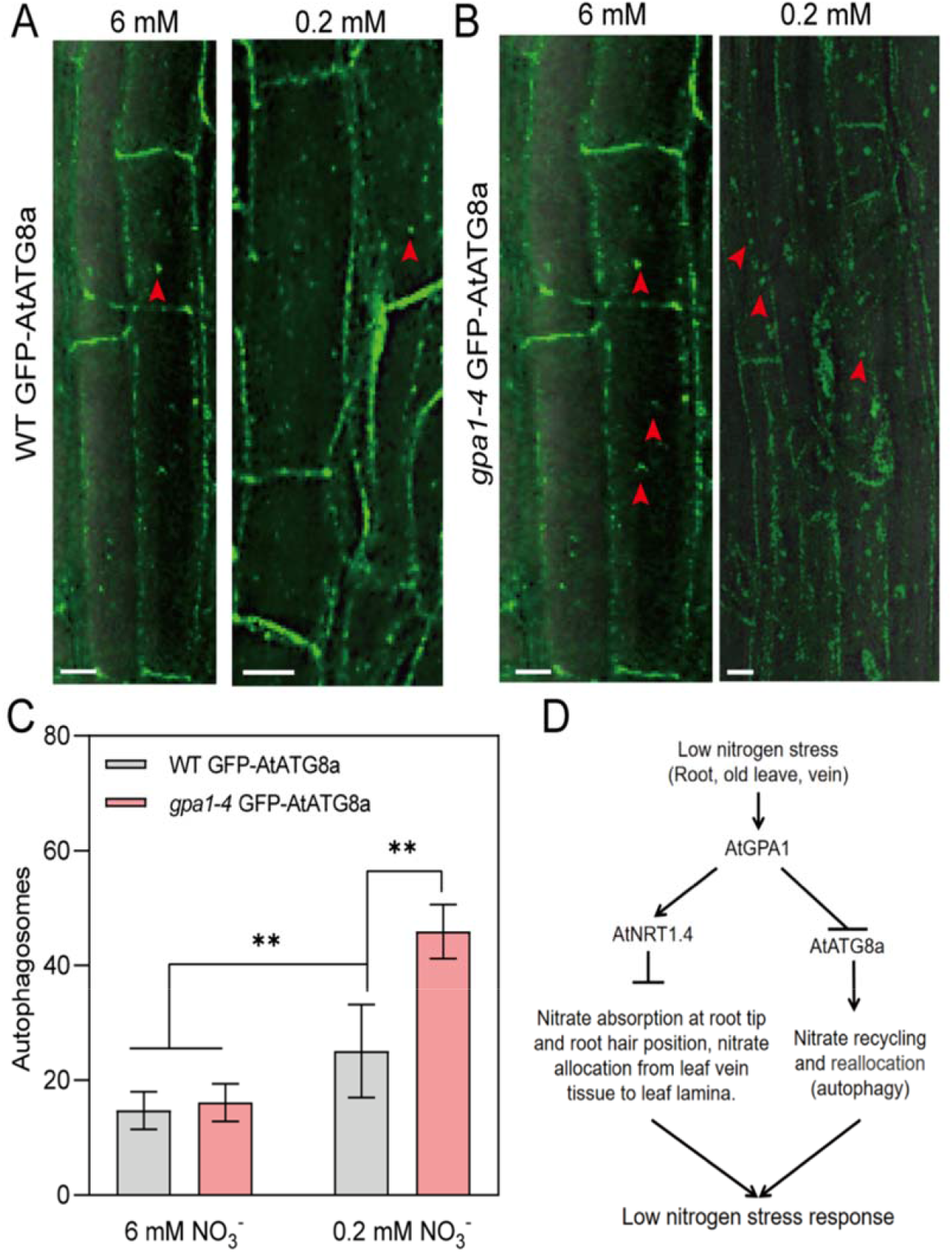
Autophagosome accumulation in the *gpa1-4* mutant and WT under normal or low-nitrate conditions. A, Autophagosomes in the root cells of seedlings of GFP-*AtATG8a* transgenic plants grown in normal-(6 mM) and low-nitrate (0.2 mM) media. B, Autophagosomes in the root cells of seedlings of *gpa1-4* GFP-AtATG8a transgenic plants on a *gpa1-4* background, grown under 6 mM and 0.2 mM nitrate condition. Red arrows indicate the autophagosomes. C, Quantification of autophagosomes in *GFP-AtATG8a* transgenic plants and *GFP-AtATG8a*/*gpa1-4* mutants. The labeled autophagosomes were counted in every root cell at the indicated times. All data are represented as mean ± SD (n = 3). Asterisks (**) above the bars represent significant differences compared to WT (*P* < 0.01 according to Student’s *t*-test). D, Regulatory model of how AtGPA1 regulates responses to low-nitrogen stress by interacting with AtNRT1.4 and AtATG8a in *Arabidopsis*.

## DISCUSSION

### AtGPA1 Regulates Responses to Low-Nitrogen Stress via AtNRT1.4 and AtATG8a

Some studies have demonstrated that G protein plays a vital role in regulating low-nitrogen stress in plants (Sun et al., 2014; Liang et al., 2018; Kaur et al., 2018; Pathak et al., 2021). However, the potential regulatory mechanism remains largely unknown. In this study, we found that the *gpa1-4* mutant was healthier than the WT under low-nitrate conditions (Fig. 1G–J), indicating that *AtGPA1* negatively regulates tolerance to low-nitrogen stress in *Arabidopsis*. We found that *AtGPA1* interacted with a nitrate transporter, *AtNRT1.4*, and a key protein, *AtATG8a*, for autophagosomes (Fig. 2). Based on these results, we constructed a model for this study to explain the underlying regulatory mechanism. *AtGPA1* is known to regulate the response to low-nitrogen stress in *Arabidopsis* via two pathways. On the one hand, the *AtGPA1*-*AtNRT1.4* interaction pair negatively regulates the absorption of nitrate from root tips and root hairs, allocation of nitrate from leaf vein tissue to leaf lamina, and plant responses to low-nitrogen stress (Fig. 7D). On the other hand, *AtGPA1* interacts with *AtATG8a* to reduce autophagosome assembly, nitrate recycling, and responses to low-nitrogen stress (Fig. 7D). In general, following receptor-mediated activation, the heterotrimeric G protein transmits external signals to cells via various downstream cytosolic effectors to regulate biological processes (Bommert et al., 2013; Urano et al., 2015; Lee et al., 2017; Roy et al., 2019). *AtGPA1* phosphorylates ABA-insensitive 1 (*ABI1*) and phospholipase Dα1 (*PLDα1*) to regulate ABA-dependent stomatal opening and closure in *Arabidopsis* (Roy et al., 2016; Hei et al., 2018; Pathak et al., 2021). The heterotrimeric G protein also interacts with plant-specific receptor-like kinases (RLKs) and receptor-like proteins (RLPs) to regulate multiple immune responses, shoot apical meristem development, and nodule formation (Bommert et al., 2013; Aranda et al., 2015; Pandey, 2020). In addition, *AtGPA1* interacts with thylakoid formation 1 (*THF1*) on the plasma membrane to regulate root development in response to exogenous D-glucose, and interacts with the cupin domain protein, *Atpirin*, to promote seed germination and early seedling development (Lapik et al., 2003; Huang et al., 2006; Xu et al., 2017). In previous studies, we have demonstrated that in *Arabidopsis, AGB1* regulates drought tolerance, morphogenesis, and the GA pathway by interacting with *AtMPK6, AtBBX21*, and *AtMYB26*, respectively (Xu et al., 2015; Xu et al., 2017; Qi et al., 2021). These results suggested that the heterotrimeric G proteins can regulate various downstream signaling pathways by interacting with various effectors. The discovery of more downstream effectors will be beneficial for elucidating novel G protein regulatory mechanisms in plants. Future studies and experiments are also needed to identify the specific biochemical mechanisms of how *AtGPA1* regulates *AtNRT1.4* and *AtATG8a*.

### *AtGPA1* Inhibits Nitrate Absorption and Allocation in Roots and Leaves by Interaction with *AtNRT1.4*

We found that *AtNRT1.4* was expressed in leaf vein tissues with sub-localization at the plasma membrane. This was similar to the findings of previous research in which *AtNRT1.4* was found to be mainly expressed in leaf petioles and leaf mid-ribs, with subcellular localization at the plasma membrane (Chiu et al., 2004; Morales et al., 2021). Interestingly, we also observed that *AtNRT1.4* was expressed extensively in roots, but not in stems (as visualized by GUS staining) (Fig. 3). Additionally, *AtNRT1.4* expression was significantly increased in response to low-nitrate conditions (Supplemental Fig. S2A). Root and leaf development were stronger in *gpa1-4* and *NRT1.4*RNAi single mutants and *NRT1.4*RNAi/*gpa1-4* double mutants than in the WT plants under low-nitrate conditions. Moreover, the phenotype of the *NRT1.4*RNAi/*gpa1-4* double mutants was closer to that of the *NRT1.4*RNAi mutants compared to the phenotype of *gpa1-4* mutants. This indicated that *AtNRT1.4* negatively regulated tolerance to low-nitrogen stress in plants, and that *AtGPA1* was an upstream factor regulating *AtNRT1.4* activity (Supplemental Fig. S3 and Fig. S4). Furthermore, the nitrate flux rate in the root hair and root tips of primary roots of seedlings was lower in the *gpa1-4, NRT1.4*RNAi, and *NRT1.4*RNAi/*gpa1-4* mutants compared to that in WT. This suggested that the *gpa1* and *nrt1.4* mutants had improved nitrate absorption from the medium to the root under normal and low-nitrate conditions (Fig. 5). A previous study showed that the disturbance function of *NRT1.5* increased nitrate distribution to root tissues, and this mutant enhanced multiple abiotic stress resistance, including salt, drought, and cadmium stress (Chen et al., 2012). Moreover, Wang et al. reported that the dysfunction of *NRT1.9* promoted plant growth by accelerating nitrate transfer to shoot from roots under high-nitrate conditions, and this mutant increased nitrate uptake indirectly (Wang et al., 2011). Our findings suggested that *AtGPA1* and *AtNRT1.4* may function in the same pathway to increase nitrate stores in vascular tissues in roots and leaf petioles. The mutation of either gene boosted nitrate absorption from the root environmental surroundings, increased nitrate release from the petiole into the leaf blade, and increased nitrate release from root vein tissue into the surrounding cells. These changes enhanced the tolerance to low nitrate stress in *gpa1-4* and *NRT1.4*RNAi mutants (Supplemental Fig. S5).

### AtGPA1 Inhibits Nitrate Recycling through *AtATG8a* under Low Nitrate Condition

We found that under low-nitrate conditions, *ATG8aOX* and *ATG8aOX*/*gpa1-4* overexpression promoted root formation and shoot growth compared to those in WT. However, there were no significant differences between mutants and the WT under normal-nitrate conditions (Supplemental Figs. S5, S6, Fig. 6). These results were in accordance with previous research, demonstrating that most ATGs are upregulated by nitrogen starvation, and that plant autophagy plays an important role in nutrient recycling under conditions of nitrogen and carbon starvation (Wang et al., 2019; Zhen et al., 2021). This was consistent with the fact that the activation of proteins in the *ATG8* family enhances tolerance to abiotic stresses, including heat, nitrogen, carbon, and drought stress (Zhou et al., 2013; Jacomin et al., 2020; Bao et al., 2020). Further analysis of autophagosome accumulation activity showed that *ATG8aOX* considerably enhanced autophagosomes in the *gpa1-4* mutant background under low nitrate conditions. We also observed that *AtGPA1* suppressed the *AtATG8a*-mediated assembly of autophagosomes (Fig. 7). Some studies have reported that the G protein-signaling pathway inhibits autophagy via inactive Gα or G protein subunits that regulate other interacting proteins in animals (Garcia et al., 2011; Liu et al., 2016; Pei et al., 2019). Plant Gα proteins are typically self-activated; for instance, RGS1 interacts with Gα and maintains the G-protein complex in its inactive state via its GAP activity (Pandey, 2019). In addition, Jiao et al. reported that *AtRGS1* increased the autophagosomes formation and autophagic flux through interaction with *AtATG8a* in response to D-glucose starvation (Jiao et al., 2019a; Jiao et al., 2019b). Combining these results, we hypothesized that *AtGPA1* inhibited nitrate recycling through *AtATG8a* under low-nitrate conditions. Phenotypic analysis indicated that *gpa1-4* and the double mutants (*NRT1.4*RNAi/*gpa1-4* and *ATG8aOX*/*gpa1-4*) were all healthier than the WT under low-nitrogen stress. Thus, we concluded that *AtGPA1* may regulate nitrate transporter and autophagy simultaneously. We suggested that *AtGPA1* mainly promoted *AtNRT1.4* as a response to low-nitrogen stress. However, further investigation is needed to determine why *AtGPA1* regulates these two pathways in response to low-nitrate stress.

## MATERIALS AND METHODS

### Plant Material and Growth Conditions

The WT was Col-0 in this study. We purchased mutant *gpa1-4* from the Arabidopsis Biological Resource Center (stock number: salk_001846). Positive mutants were screened using the triple primer method, and primers were searched on T-DNA Primer Design by stock number (http://signal.salk.edu/tdnaprimers.2.html). We obtained *NRT1.4*RNAi mutant lines using the vacuum infiltration method, and the interference was successful in *NRT1.4*RNAi transgenic lines #12, #14, and #38. The *NRT1.4*RNAi*/gpa1-4* double mutant lines #22 and #25 were the hybrid offspring of *NRT1.4*RNAi #12 and *gpa1-4*, whereas *NRT1.4*RNAi*/gpa1-4* double mutant line #32 consisted of the hybrid offspring of *NRT1.4*RNAi #38 and *gpa1-4*. We obtained *ATG8a* overexpression lines by transforming the pCAMBIA1302:*ATG8a* vector into the WT. The *ATG8aOX/gpa1-4* double mutant lines #3 and #6 were hybrid offspring of *ATG8aOX* #3 and *gpa1-4*, and the *ATG8aOX/gpa1-4* double mutant line #17 consisted of hybrid offspring of *ATG8aOX* #7 and *gpa1-4*.

We sterilized the seeds of WT and transgenic plants with 70% ethanol and washed them with distilled water; subsequently, the sterilized seeds were washed again with 1% sodium hypochlorite solution in a clean laboratory. Sterilized seeds were spread on Murashige and Skoog (MS) medium, and refrigerated at 4 °C for 3 d of stratification. During this period, the seedlings were transferred to normal or low-nitrogen medium for 7 d. Finally, 10-d-old seedlings were photographed and the total root length, total root surface, fresh weight, and chlorophyll content were measured. The MS or nitrogen medium was cultured in a culture room at 30 °C and a photoperiod of 16 h.

### Yeast Two-Hybrid

Briefly, we constructed a fusion expression vector, pDHB1-AtGPA1, by inserting the coding region of *AtGPA1* into the bait vectors pDHB1, and the prey vectors pPR3N-AtNRT1.4 or pPR3N-AtATG8a were obtained from pPR3N. All primers mentioned in Table S1 (Supplemental Table S1) were used for vector construction. Empty bait or prey vectors were transformed together with pDHB1-AtGPA1 or pPR3N-AtNRT1.4 plasmids as negative controls. Interactions of pDHB1-APP and pPR3N-Fe65 were used as positive controls. We co-transformed vectors into NMY51 yeast cells, which are used for membrane interaction proteins according to the lithium acetate method.

### AtNRT1.4 Pull-Down by GST-AtGPA1 Agarose Beads

The coding region of *AtGPA1* and *AtNRT1.4* were constructed into the vector pGEX-4T-1 and pCold, respectively. The fusion expression vector His-AtNRT1.4 or GST-AtGPA1 were expressed in *Escherichia coli*, added IPTG to induce the fusion protein, and the target proteins were purified using Ni or glutathione agarose beads, respectively, and the protein samples were analyzed by sodium dodecyl sulfate-polyacrylamide agarose gel electrolysis.

### Bimolecular Fluorescence Complementation (BiFC) Assay

The coding region of *AtGPA1* was cloned into the pSPYCE vector. The coding region of *AtNRT1.4* or *AtATG8a* was separately cloned into the pSPYNE vector, and the fusion expression vectors were co-transformed into protoplasts of *Arabidopsis* mesophyll. After 12 h, we captured yellow fluorescent protein (YFP) fluorescence and chloroplast autofluorescence signals using a laser scanning confocal microscope. YFP fluorescence signals were measured in the 500–570-nm wavelength range, and chloroplast autofluorescence was measured in the 630–700-nm wavelength range.

### GUS Staining Analyses

To obtain *ProAtGPA1-GUS* and *ProAtNRT1.4-GUS* seedlings, we inserted the promoter of *AtGPA1* and *AtNRT1.4* into the PBI121 plasmid, and then transformed it into the WT. Homozygote seedlings were grown on MS medium, and the seedlings were treated with absolute ethanol (a destaining solution) to remove chlorophyll. The whole plants were then stained with GUS staining buffer under dark conditions for 12 h at 28 °C. The pictures of GUS seedlings were observed and captured using a light microscope (SZX16, Olympus) with a digital camera (DP-73, Olympus).

### Subcellular Localization of *AtGPA1* and *AtNRT1.4*

The coding regions of *AtGPA1* and *AtNRT1.4* were inserted into 16318hGFP to construct the fusion expression vector. The amplification primers were 16318hGFP-AtGPA1-F/R and 16318hGFP-AtNRT1.4-F/R (Supplemental Table S1). The empty 16318hGFP vector was used as a positive control. We observed GFP signals using confocal microscopy using a Leica (Yoo et al., 2007).

### Measurement of the Net Nitrate Flux

We tested the net fluxes of nitrate using the SIET method as described previously (Zheng et al., 2013). The net fluxes of nitrate in the root tip and root hair zone were measured, and each value represented the means of the three plants. We performed this experiment at Xuyue Science and Technology (Beijing, China).

### Measurement of Root Traits

After cultivation on a normal or low-nitrogen medium for 7 d, we measured the total root area and total root length using Spectra Scan-R. All measurements were performed in at least six replicates.

### Measurement of Fresh Weight and Chlorophyll Content

For each material, three seedlings were weighed for one “fresh weight” data using at least three biological replicates. The leaves of seedlings were soaked in overnight in 80% acetone while being swayed at 100 rpm in the dark. We measured the absorbance of the extracting solution at 645 nm and 663 nm using Thermo Scientific Microplate Reader (Lichtenthaler, 1983).

### Observation of Autophagosomes

To observe the autophagosomes in the root cells of seedlings, *gpa1-4* mutant and WT were grown under normal or low nitrate conditions and the seedling roots were treated with 1 μM concanamycin A (CA) for 12 h. We monitored GFP fluorescence signal at 505–550 nm and photographed the signals for each material. Further, we calculated GFP fluorescence points, which represented the number of autophagosomes with ImageJ software.

### Statistical Analysis

Data of all experimental analyses were obtained from at least three biological replicates. The means and standard deviations of all the values were analyzed based on one-way analysis of variance, and Duncan’s multiple range test at a significance level of *P* < 0.05 using SPSS software version 21 (International Business Machines Corp, New York, USA).

## Accession Numbers

The following were the accession numbers of the gene sequences: *AtGPA1* (AT2G26300), *AtNRT1.4* (AT2G26690), and *AtATG8a* (AT4G21980), in the *Arabidopsis* TAIR10 website (http://www.arabidopsis.org/).

## SUPPLEMENTAL DATA

**Figure 1. Phenotype of *gpa1-4* mutants and WT under low nitrate conditions**.

**Figure 2. *AtGPA1* interacts with *AtNRT1.4* and *AtATG8a***.

**Figure 3. Tissue-specific expression and subcellular localization of *AtGPA1* and *AtNRT1.4***.

**Figure 4. Phenotype of single mutants, *gpa1-4* and *NRT1.4*RNAi, and double mutant, *NRT1.4*RNAi/*gpa1-4*, under low-nitrogen conditions**.

**Figure 5. Nitrate efflux rate of WT and different mutants in the root tip and root hair zones under normal or low-nitrate conditions**.

**Figure 6. *AtATG8a* overexpression in WT and *gpa1-4* backgrounds increases tolerance to low nitrate stress**.

**Figure 7. Autophagosome accumulation in the *gpa1-4* mutant and the WT under normal conditions and low-nitrate conditions**.

**Figure S1. Mutant *gpa1-4* shows better growth and has wider leaves than the WT under low nitrate conditions**.

**Figure S2. Expression analysis of *AtGPA1, AtNRT1.4*, and *AtATG8a* in different mutant or transgenic plants under low-nitrogen conditions**.

**Figure S3. Phenotypes of WT plants and *NRT1.4*RNAi lines grown under normal or low-nitrogen conditions**.

**Figure S4. Tolerance to low nitrate stress is enhanced in the double mutant, *NRT1.4*RNAi*/gpa1-4***.

**Figure S5. Phenotype of *AtATG8a*-overexpressing plants on a WT background under low nitrate stress. Figure S6. Phenotype of the double mutant, *ATG8aOX/gpa1-4*, under low nitrate conditions**.

**Supplemental Figure S1**. Mutant *gpa1-4* shows better growth and has wider leaves than the WT under low nitrate conditions.

**Supplemental Figure S2**. Expression analysis of *AtGPA1, AtNRT1.4*, and *AtATG8a* in low-nitrogen conditions or in different mutant or transgenic plants.

**Supplemental Figure S3**. Phenotype of *NRT1.4*RNAi lines on a background of WT or *gpa1-4* mutants under normal or low-nitrogen conditions.

**Supplemental Figure S4**. Tolerance to low nitrate stress is enhanced in the double mutant, *NRT1.4*RNAi*/gpa1*.

**Supplemental Figure S5**. Phenotype of *ATG8aOX* lines.

**Supplemental Figure S6**. Phenotype of *ATG8aOX/gpa1-4* lines.

**Supplemental Table S1**. Primers and ingredients of the nitrate medium used in this study.

## ACKNOWLEDGEMENTS

The authors are grateful to Prof Pierre Delaplace, University of Liege, for revising this manuscript. This work was supported by the Agricultural Science and Technology Innovation and China Scholarship Council (contract no. 202003250115). We would like to thank Editage (www.editage.cn) for English language editing.

## Notes

1 This work is supported by the Agricultural Science and Technology Innovation and the scholarship from the China Scholarship Council (NO. 202003250115).

